# Extracting and Updating Compact Colored *k*-mer Indexes

**DOI:** 10.64898/2026.11.06.764792

**Authors:** Lore Depuydt, Jarno N. Alanko, Camille Marchet, Simon J. Puglisi

**Affiliations:** Univ. Lille, CNRS, Centrale Lille, UMR 9189 CRIStAL, F-59000 Lille, France; Department of Computer Science, University of Helsinki, Finland

**Keywords:** colored *k*-mer index, spectral Burrows-Wheeler transform, colored de Bruijn graph, set operations

## Abstract

Colored *k*-mer indexes record, for every *k*-mer of a collection of genomes, the genomes or species that contain it, and underpin pseudoalignment and sequence search against large collections. Construction of such an index is expensive, yet the collection it represents is rarely final: genomes are retracted or found to be contaminated, users need an index of a single species or clade, and new genomes keep arriving. Rebuilding the index after every such change requires the original sequences and repeats the costly construction. We show that colored indexes based on the spectral Burrows-Wheeler transform (SBWT) can instead be updated efficiently in batches, without the original sequences, and with a result equivalent to the index built from scratch. Given an SBWT and any subset of its *k*-mers, we first extract the SBWT of this subset directly from the input, in one linear pass and without building a new index. Removing genomes, intersection and difference all reduce to this operation. We then carry the colors through every operation, by finding from the inputs alone where the color sets of the result change, so that removing or extracting genomes, as well as the union, intersection and difference of two colored indexes, each directly yield a complete colored index. In our experiments, editing paid off for large results: removing a genome from an index of 3 682 *E. coli* genomes took 54 seconds instead of 13 minutes for a rebuild, and adding one took about a minute. On AllTheBacteria, with 192 billion *k*-mers and 12 733 species, extracting *M. tuberculosis*, with 180 094 genomes, took less than four hours, and set operations then traced contamination of its color, some of which the next release of AllTheBacteria still contains. Our work turns large colored *k*-mer indexes from static snapshots of a collection into resources that can be maintained as the collection evolves. Our implementation is available at https://github.com/LoreDepuydt/colored-sbwt-setops.

## 1 Introduction

A colored *k*-mer index records, for every *k*-mer of a collection of genomes, which genomes, or *colors*, contain it. Such indexes underpin pseudoalignment [1, 2, 3, 4] and sequence search against collections that now count up to millions of bacterial genomes, such as the 661K collection [5] and AllTheBacteria [6]. State-of-the-art indexing approaches fall into two families: Themisto [2] and MetaGraph [4] employ variants of the Burrows-Wheeler transform (BWT), whereas Fulgor [3], Bifrost [7] and more recently also MetaGraph make use of minimizer hashing. In this work, we focus on the Burrows-Wheeler family of indexes, namely the spectral Burrows-Wheeler transform (SBWT) [8] used by Themisto.

Constructing such an index is expensive: it can take hours or even days and hundreds of GiB of memory. Yet, the collection it represents is rarely final. New genomes keep arriving, while existing assemblies turn out to be contaminated or misassigned, or are retracted altogether. Users may also want an index of only a subset of the collection, such as all genomes of a single species or clade, which is smaller and faster to query. Finally, when a collection is re-released, users want to know which *k*-mers the new release adds and which it drops. Each of these tasks currently requires reindexing from the sequences, which may not be at hand: the SBWT of the 661K collection takes 52 GiB, against 2.57 Tbp of sequence.

In this paper, we ask whether such changes can be made directly on the index, without the original sequences, and faster than a rebuild when the result is large. The result should be equivalent to the index built from scratch: it has the same SBWT, and every *k*-mer has the same color set (the set of colors containing it). In our earlier work [9], we computed the union, intersection and difference of two uncolored SBWTs from a *merge plan* of the two inputs. Two things were missing: colors, and an efficient way to repair the index when *k*-mers disappear (Figure 1). Repair is needed because, besides the *k*-mers, an SBWT stores *dummy nodes* ($-padded prefixes that give every *k*-mer a predecessor): deleting *k*-mers leaves some kept *k*-mers without a predecessor and some dummy nodes leading nowhere.

**Figure 1.**
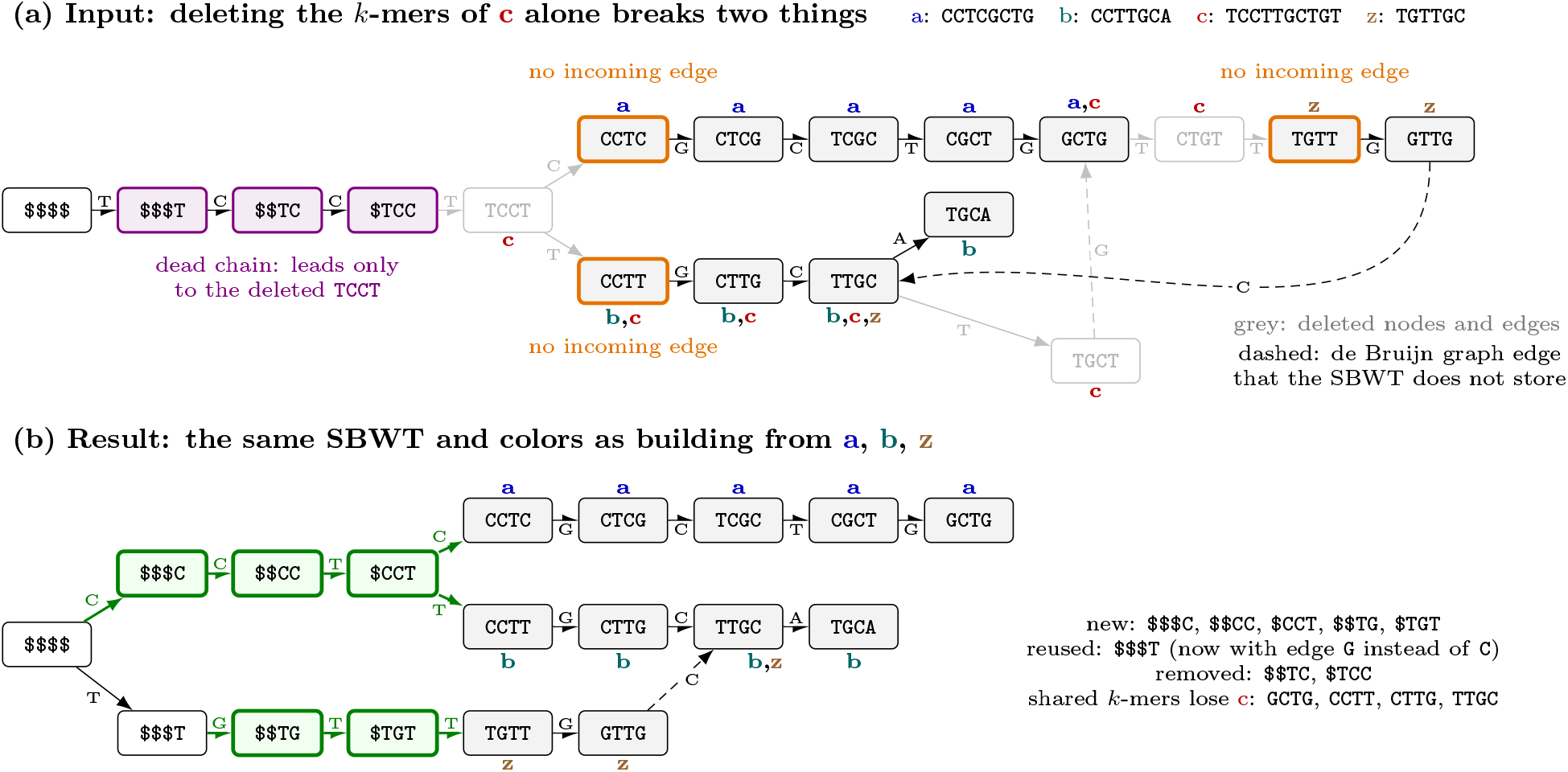
The running example as SBWT graphs (*k* = 4). The SBWT keeps one incoming edge per node. Dashed edges are de Bruijn graph edges it does not store (with them, every genome is a path). (a) The input, with the color set of each *k*-mer written next to it. Deleting the three *k*-mers of c alone and their edges (grey) leaves three kept *k*-mers without an incoming edge (orange) and a dummy chain leading nowhere (violet). (b) The result: five new dummy nodes (green), one reused, two removed. The shared *k*-mers stay, without c.

We show that removing and extracting genomes, intersection, difference and union work on the colored index, and we implemented them in Rust^1^, extending the sbwt library^2^. Our contributions are as follows.

Given an SBWT and any subset *K* of its *k*-mers, we extract the SBWT of *K* directly from the input (Section 3.1), without the original sequences and without building a new index. In one linear pass, we insert each missing dummy node into the colex order of the input, at a position found by a forward search, and drop those leading nowhere. In our experiments, this repaired the SBWT 4 to 126 times faster than our earlier work (Section 4.2). Given the input’s group starts, our extraction takes *O* (*n* + |*Q* |*k*) time, where *n* is the size of the input and *Q* the result nodes that lost their in-neighbor. Removing genomes, extracting a species, intersection and difference are all such extractions.

We carry the colors through every operation (Section 3.2). A colored index such as that of Themisto stores every distinct color set once, and attaches one to the *k*-mers where the color set changes or a unitig ends, which the result of an edit must determine anew. We observe that the color changes of the result can be found from the inputs alone: the color set of a *k*-mer in the result depends only on its color sets in the inputs, so every place where it changes in the result shows up in the inputs as well, and one walk over the unitigs of the inputs finds every such place. Every operation thus directly yields a complete colored index, without the genomes and without computing the color set of every *k*-mer.

We show that editing pays off when the result is large (Section 4.1): removing a genome from an index of 3 682 *E. coli* genomes took 54 s instead of 13 min for a rebuild from the genomes (14*×*). Adding a genome to the *E. coli* index took about a minute, including building the index of the new genome, instead of 13 min for a rebuild (13*×*). Only for small results, such as extracting a few percent of the genomes, is a rebuild faster, and an edit holds its whole input index in memory.

In a case study on AllTheBacteria (Section 4.3), we show that editing can also answer questions that a rebuild cannot. Extracting *M. tuberculosis* from a species-colored AllTheBacteria index, with 192 billion *k*-mers, took less than four hours, and kept, for every *k*-mer of the species, all other species that share it. With set operations, one heuristic filter and no assemblies, we then removed about a third of the *M. tuberculosis* color as likely foreign, including contamination that the next release still contains.

### 1.1 Related work

Set operations on *k*-mer sets are supported by *k*-mer counters such as KMC [10], by the dynamic representation CBL [11], and by masked superstrings [12]. For compressed indexes, the usual way to combine two collections is to merge them, as for BWTs [13, 14, 15, 16], Wheeler graphs [17], the SBWT [9] and BOSS [18, 17], a de Bruijn graph representation [19] close to the SBWT. Merging only adds *k*-mers, so no *k*-mer loses its predecessor and no dummy node has to be added; removing *k*-mers is the harder case.

The alternative is a dynamic index, which is updated in place. Dynamic compressed text indexes insert and delete strings [20, 21, 22], dynamic de Bruijn graphs insert and delete *k*-mers [23, 24, 25], and those based on BOSS repair their dummy nodes one deleted *k*-mer at a time [24] or for a batch with an *O*(*nk*) merge [25]; the graph BWT can remove haplotypes [26]. Most update one element at a time, paying in space, query time or reorganization [25], whereas we apply a batch in one linear pass and get the static index.

Most colored *k*-mer indexes are static [2, 3, 27, 4], and recent work compresses their color sets ever further by exploiting the redundancy between them [28, 29], which makes them no easier to update in place. Some can add genomes, by merging [18], with the Bentley-Saxe transformation [30], or by insertion [31]. Approximate indexes based on Bloom filters [32, 33, 34, 35, 36] are easier to extend, as the dynamic hierarchical interleaved Bloom filter shows [37], but answer queries with false positives. Removal is rarer still: Bifrost supports it through its API [7], and MetaGraph can keep its graph in a dynamic representation [4], but both then use a dynamic structure, larger and slower to query than a static compressed index. To our knowledge, no method removes genomes from, or extracts a subset of, a static compressed colored index with a result equivalent to construction.

## 2 Preliminaries

Since our operations edit an index without the sequences it was built from, this section recalls what the index stores, and the properties of the SBWT that make editing it possible. We follow the definitions of the SBWT paper [8] and of our earlier work [9], and illustrate each of them on a running example.

The example has *k* = 4 and, for readability, uses the forward strand only. It consists of three related strains, a: CCTCGCTG, b: `CCTTGCA` and c: TCCTTGCTGT, and an unrelated genome z: TGTTGC. Throughout the paper, our goal is to remove the c strain from the index. A real index covers both strands: every genome is added together with its reverse complement, under the same color, so a *k*-mer and its reverse complement always have the same color set. Our operations do not use this pairing but preserve it: a *k*-mer and its reverse complement are kept or removed together, so the result is the index built from scratch on both strands.

### SBWT

Strings are over the DNA alphabet Σ = *{*A, C, G, T*}* , extended with a special character $ ∉ Σ that is smaller than all characters of Σ. The colexicographic (colex) order compares strings from right to left. The SBWT of a set *K* of *k*-mers represents a graph in which every *k*-mer has a predecessor. Some *k*-mers have no predecessor in *K*, so the SBWT adds $-padded *dummy nodes*, which lead to them from the root $^*k*^. Concretely, the padded *k*-spectrum *K*^+^ consists of *K*, the root, and, for every *k*-mer *c*_1_ *· · · c*_*k*_ *∈ K* such that *c*_1_*· · · · · ·c*_*k−*1_ is not a suffix of any *k*-mer in *K*, the dummy chain $^*k−*1^*c*_1_, $^*k−*2^*c*_1_*c*_2_, … , $*c*_1_ *c*_*k−*1_. In the example, TCCT is the only *k*-mer without a predecessor, so the only dummy chain is $$$T, $$TC, $TCC.

The SBWT graph is a subgraph of the node-centric de Bruijn graph. The latter has an edge from *x* to *y* labeled *c* whenever *y* = *x*[2..*k*]*c*. To give every node other than the root a single parent, the SBWT graph keeps only the incoming edge from its smallest in-neighbor in colex order. For example, TTGC has the two in-neighbors `CTTG` and `GTTG`, and the SBWT graph only keeps the edge from `CTTG`. The SBWT graph is a tree rooted at $^*k*^ when every *k*-mer can be reached from the root, and otherwise also contains cycles of *k*-mers, such as `AC` and `CA` for the string ACA and *k* = 2. The SBWT lists the nodes of *K*^+^ in colex order, numbered from 0 to *n−*1, and stores for every node the labels of its outgoing edges in this graph (Table 1), as a bit matrix with a row per node and a column per character.

**Table 1.**
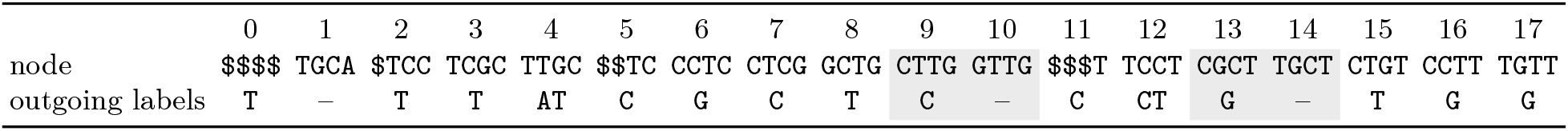
The SBWT of the running example: its nodes in colex order, with the labels of their outgoing edges in the SBWT graph. Shaded: the two suffix groups with more than one node. The *k*-mers are shown but not stored.

The nodes that share their last *k−*1 characters are consecutive in colex order, and form a *suffix group*, or simply a *group*. All nodes of a group have the same out-neighbors in the de Bruijn graph. The first node is the smallest in-neighbor of each of them in colex order, so only this node stores the outgoing edges of the group. For example, `CTTG` and `GTTG` form a group, and the edge to `TTGC` is stored only at `CTTG`. Our operations traverse the de Bruijn graph, and thus need the *group starts*, a bit vector of *n* bits that marks the first node of every group. The group starts are not visible in the rows, and take *O*(*nk*) time to compute [9].

Two properties of the colex order turn SBWT graph traversal and search into rank and select queries. First, the *i*-th node with an outgoing edge labeled *c*, in colex order, leads to the *i*-th node that ends with *c*. In Table 1, for example, the edges labeled C leave nodes 5, 7, 9, 11 and 12, and lead to nodes 2 to 6, in the same order. With *C*[*c*] the position of the first node that ends with *c*, and rank_*c*_(*p*) the number of nodes before *p* with an outgoing label *c*, the outgoing edge labeled *c* of node *p* therefore leads to node LF(*p, c*) = *C*[*c*] + rank_*c*_(*p*). Conversely, following the incoming edge of a node backward takes a single select query on the column of its last character. Second, the same counting applies to all ranges of nodes. For a string *α* with |*α*| *< k*, the nodes that end with *α* are consecutive in colex order, and if they form the range [*b, e*), the nodes that end with *αc* form the range [LF(*b, c*), LF(*e, c*)), since their incoming edges all come from [*b, e*). This *forward search* finds the range of *α* in |*α*| steps, starting from the range [0, *n*) of all nodes.

### Colored indexes

The SBWT only represents a set of *k*-mers. A colored *k*-mer index also records, for every *k*-mer *x* of a collection of genomes, its *color set* col(*x*), the set of *colors* that contain *x*. A color can be a genome, or a group of genomes such as a species. In the example (Figure 1a), col(`TTGC`) = *{b, c, z}*. Dummy nodes are not *k*-mers of any genome, and have no color set. As in Themisto [2], we represent a colored index by the SBWT, the distinct color sets, and, for a subset of *k*-mers that we call *key k-mers*, the identifier of their color set. The color set of any other *k*-mer is found by walking forward along its *unitig* up to the next key *k*-mer. A unitig is a maximal path in the de Bruijn graph of *K* (not in the SBWT graph, in which every node has in-degree one) such that, for every two consecutive nodes *x* and *y, x* has out-degree one and *y* has in-degree one. Since the walk must stop at the end of its unitig, the last *k*-mer of every unitig is a key *k*-mer (on a cyclic unitig, we pick one *k*-mer as its last). In the example, these are CGCT, CTTG, GTTG, TCCT, TGCA, TGCT and `TTGC`. Themisto also stores the color set identifier of every *d*-th *k*-mer along a unitig, counted back from its last *k*-mer, the *checkpoint k*-mers, so that a look-up takes at most *d* steps.

Beyond the topology, the color set can also change along a unitig, which thus splits into *colored subunitigs*, the maximal subpaths whose *k*-mers all have the same color set. For example, the unitig GCTG, CTGT, TGTT,GTTG consists of the colored subunitigs `GCTG` with *{a, c }*, `CTGT` with *{c}* , and TGTT, GTTG with *{z}*. For the walk to return the right color set, the last *k*-mer of every colored subunitig must also be a key *k*-mer. Here, this adds `GCTG` and `CTGT`. The operations in this paper *walk the unitigs* of an index with the SBWT, in *O*(*n*) time given the group starts, which give the in- and out-degree of every node in constant time. The same walk also splits every unitig into its colored subunitigs.

### Merge plan

An operation on two indexes, such as a union, must also relate the nodes of one index to those of the other. Given two SBWTs with padded *k*-spectra *A*^+^ and *B*^+^, the merge plan [9] describes the colex order of *A*^+^*∪ B*^+^. We store it as two bit vectors that mark, for every position, whether its *k*-mer is in *A*^+^ and in *B*^+^, and a third that marks the first position of every group. The merge plan is computed once per operation, in *O*(*nk*) time. For operations on two inputs, *n* denotes the number of positions of the merge plan, |*A*^+^ *∪ B*^+^|.

## 3 Methods

Section 3.1 shows how to extract the SBWT of any subset of the *k*-mers of an SBWT, which gives the SBWT of the result of every operation except the union, whose result is not a subset of either input. As no *k*-mer loses its predecessor in a union, its SBWT follows directly from the merge plan [9]. Section 3.2 then carries the colors through every operation.

### 3.1 Extraction: removing *k*-mers from an SBWT

Let *K* be a subset of the *k*-mers of an SBWT. Computing the SBWT of *K* from the input SBWT, without the original sequences and without building a new index, is what we call an *extraction*. Removing genomes is an *extraction by colors*, in which *K* consists of the *k*-mers that keep at least one remaining color. The intersection *A ∩ B* and the difference *A \ B* are extractions from *A*, in which *K* consists of the *k*-mers of *A* that are, or are not, in *B*, as marked in the merge plan of *A* and *B* (Section 2). Deleting the *k*-mers that are not in *K* is straightforward, but the result must also contain exactly the dummy nodes of *K*^+^, on which the SBWT of *K* is defined (Section 2). After the deletion, the remaining SBWT graph has both too many and too few dummy nodes: some dummy chains of the input now lead only to deleted *k*-mers, and some kept *k*-mers, the *sources*, lose all their in-neighbors and need a new dummy chain. Figure 1 shows both defects on the running example. Removing strain c deletes the three *k*-mers that occur in c alone, TCCT, CTGT and `TGCT`. The kept *k*-mers `CCTC` and `CCTT` were reached through `TCCT`, and `TGTT` through `CTGT`, so these three are sources. Conversely, the dummy chain $$$T, $$TC, $TCC existed only to reach TCCT, and now leads nowhere. Repairing both defects must give the SBWT of Figure 1b, the one built from a, b and z directly.

Where our earlier work [9] built the missing dummy nodes as a separate SBWT and merged it in with a three-way merge plan (the *three-way merge*), our main observation is that they can be *inserted* directly into the colex order of the input, at positions that a forward search finds.

#### 3.3.1. Finding the defects in one pass

The sources are found with the incoming-edge coverage pass of our earlier work [9], in *O*(*n*) time. This pass records, for every node, whether it has a kept in-neighbor. The kept nodes other than the root that have none are the sources, which need a new dummy chain.

The same pass also finds the *dead ends*: the kept dummy nodes whose group keeps no outgoing edge. A *dead chain* consists of a dead end and of its ancestors other than the root. We walk up each dead chain from its dead end, one select query per step, and mark its nodes as dead. A walk stops at a node that is already marked dead, so every node is marked at most once. Since the walk does not inspect the subtrees of these nodes, it may also mark nodes as dead that still lead to kept nodes, which have then lost their only in-neighbor. A final sweep over the outgoing edges of the dead nodes adds these kept nodes to the sources. Marking a node dead that still leads to a kept node is safe: it lies on the dummy chain of the child that became a source, and the repair unmarks it again (Lemma 1). We denote the set of sources by *Q*. In the example, the only dead end is $`TCC`, whose only edge led to TCCT (Figure 1a), and the walk marks its chain $`TCC`, $$TC, $$$T as dead. The sources are *Q* = *{*CCTC, CCTT, TGTT *}*, and since $$$T also starts the dummy chain of TGTT, the repair eventually unmarks it again. Since every node is marked dead at most once and the SBWT graph has fewer than *n* edges, finding both defects takes *O*(*n*) time in total, and three bit vectors of *n* bits.

##### Algorithm 1

Inserting the dummy chains of the sources into the colex order of the input

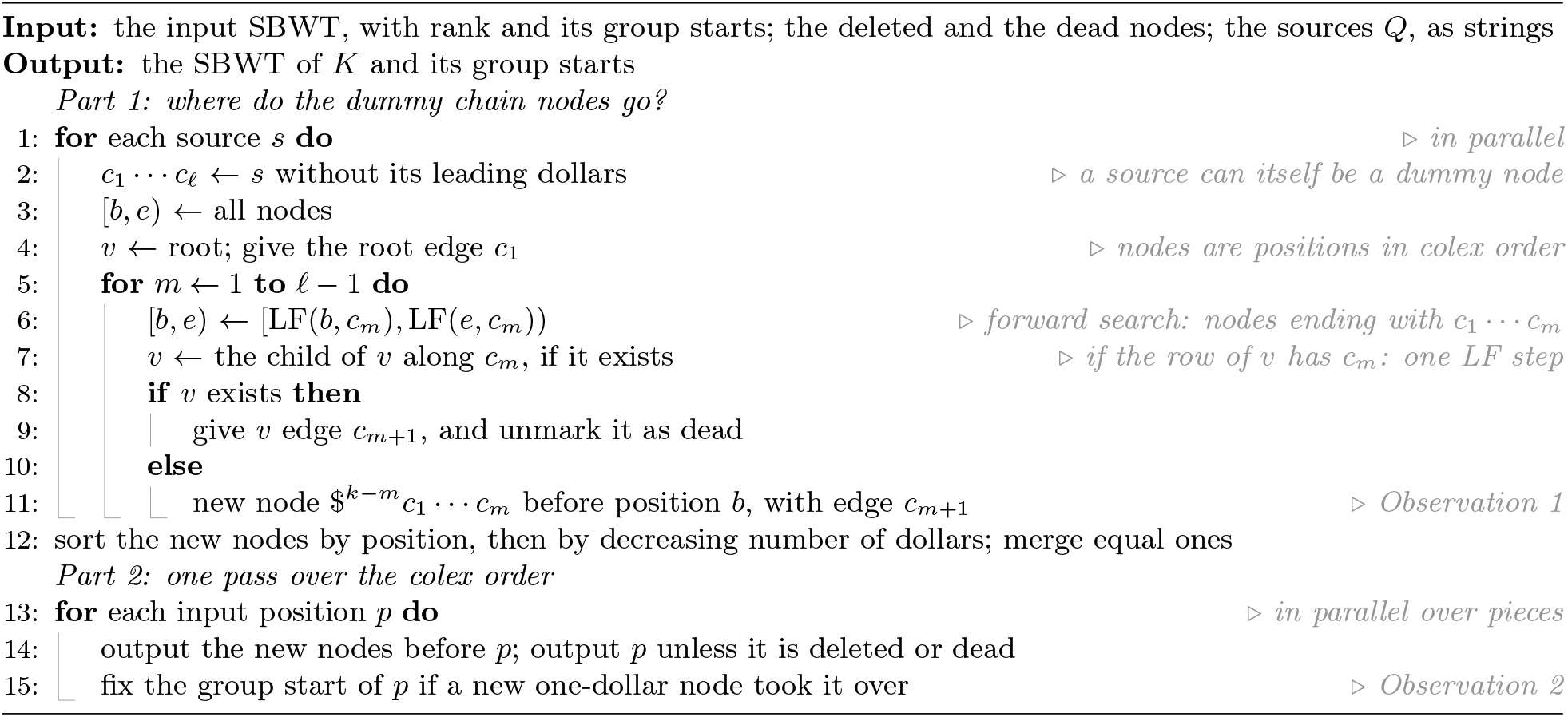

#### 3.1.2 Inserting the missing dummy nodes, removing deleted and dead nodes

It remains to give every source its dummy chain. A source *s* = $^*k−l*^*c*_1_ *· · · c*_*l*_, with *l* = *k* unless the source is itself a dummy node, needs the dummy chain $^*k−*1^*c*_1_, $^*k−*2^*c*_1_*c*_2_, … , $^*k−l*+1^*c*_1_*· · · c*_*l−*1_. Some of these nodes may already exist in the input, possibly marked dead, and the others must be added. We insert each of them directly into the colex order of the input (*insertion*), instead of the three-way merge. For each new node $^*k−m*^*c*_1_*· · · c*_*m*_, this requires knowing where it goes in the colex order, and which group it belongs to, so that the rows of the SBWT remain correct. Both questions are answered by a forward search of *c*_1_*· · · c*_*m*_ in the input. Its range is never empty: every node of the *input* other than the root has an in-neighbor, so the source has an ancestor in the input, possibly deleted, that ends with *c*_1_ *· · · c*_*m*_.

##### Observation 1

*The new node* $^*k−m*^*c*_1_*· · · · · ·c*_*m*_ *comes right before the first input node that ends with c*_1_ *· · ·c*_*m*_. *When several new nodes come before the same node, they are ordered by decreasing number of dollars*.

All nodes ending with *c*_1_ *c*_*m*_ agree with the new node on these *m* characters, and the new node next has a $, which is smaller than every character, so it precedes all of them. Every node before the first of them is smaller on its last *m* characters, and thus also precedes the new node. Two new nodes before the same node end with two suffixes of that node, one of which is a suffix of the other, so the node with more dollars has a $ where the other has a character, and comes first. For example, $$TG goes right before `GCTG`, the first of the three input nodes that end with `TG`, and $$$`C` and $$`CC` both go before $`TCC`, in this order.

##### Observation 2

*A new node with one dollar becomes the first node of the group of the next node, and one with two or more dollars forms its own group. Thus the groups, and hence the rows, remain correct*.

The new node $*c*_1_ *· · · c*_*k−*1_ has the same last *k −* 1 characters as the node after it, which is the first node that ends with *c*_1_*· · · c*_*k−*1_. With two or more dollars, the last *k−* 1 characters of the new node start with a $, and no other node shares them. For example, $`CCT` takes over the group of the deleted `TCCT`.

Algorithm 1 puts the two observations together, with the sources spelled out as strings as in our earlier work [9]. In Part 1, the dummy chain of each source is followed from the root while its nodes exist, and new nodes are created from the first missing one on. The existing nodes of the dummy chain are unmarked as dead, and nodes shared by several sources are merged, with all their edges. Part 2 outputs the new nodes in place, drops the deleted and the dead nodes, and writes the rows of the result: every group gets the edges of the input that lead to a node of the result, and the edges of the new dummy chains.

*Example*. Table 2 shows the result, with the positions of the input in Table 1. The five new dummy nodes stand where Observations 1 and 2 put them, and $`CCT`, which the sources `CCTC` and `CCTT` share, gets both edges `C` and `T`. The source `TGTT` needs $$$`T`, which exists and is unmarked as dead again. Part 2 leaves out the deleted positions 12, 14 and 15, and the dead positions 2 and 5, which no source needs. This gives exactly the SBWT built from a, b and z.

**Table 2.**
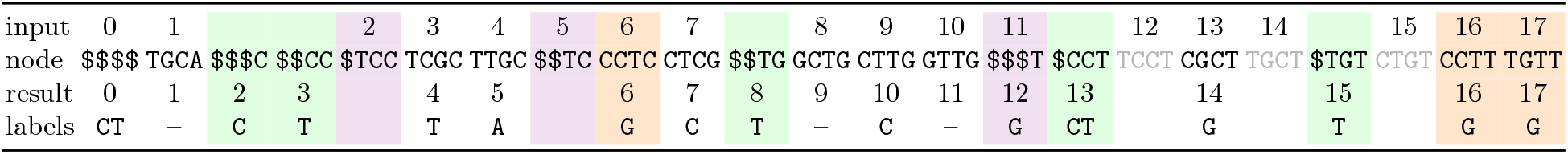
The SBWT after removing strain c: the nodes of the input in colex order with the five new dummy nodes inserted, their positions in the input and in the result, and the labels of their outgoing edges in the result. Green: new dummy nodes; violet: marked dead nodes; orange: sources; grey: deleted *k*-mers.

| input | 0 | 1 |  |  | 2 | 3 | 4 | 5 | 6 | 7 |  | 8 | 9 | 10 | 11 |  | 12 | 13 | 14 |  | 15 | 16 | 17 |
|---|---|---|---|---|---|---|---|---|---|---|---|---|---|---|---|---|---|---|---|---|---|---|---|
| node | \$\$\$\$ | TGCA | \$\$\$C | \$\$CC | \$TCC | TCGC | TTGC | \$\$TC | CCTC | CTCG | \$\$TG | GCTG | CTTG | GTTG | \$\$\$T | \$CCT | TCCT | CGCT | TGCT | \$TGT | CTGT | CCTT | TGTT |
| result | 0 | 1 | 2 | 3 |  | 4 | 5 |  | 6 | 7 | 8 | 9 | 10 | 11 | 12 | 13 |  | 14 |  | 15 |  | 16 | 17 |
| labels | CT | – | C | T |  | T | A |  | G | C | T | – | C | – | G | CT |  | G |  | T |  | G | G |

##### Lemma 1

*The extraction outputs the SBWT of K and its group starts*.

*Proof sketch*. The output contains the *k*-mers of *K* and the root, so it remains to show that its dummy nodes are those of *K*^+^. A kept *k*-mer needs a dummy chain in *K*^+^ exactly when it has no in-neighbor in *K*: either its in-neighbors in the input were all deleted, and it is a source, or it had none in the input either, and its dummy chain is in the input. Algorithm 1 keeps or inserts every node of the dummy chain of a source. A node of a needed input dummy chain that a walk marked dead leads to a kept node, which the sweep turns into a source, so it lies on the dummy chain of that source and is kept as well. Conversely, an input dummy node that no kept *k*-mer needs leads only to deleted *k*-mers, so it lies on the walk up from a dead end, and is left out. Observations 1 and 2 put every new node at its colex position and in its group, and every group keeps the edges of the input that lead to a node of the result and gains those of the new dummy chains, which are exactly the edges of the SBWT graph of *K*^+^.

The extraction sorts no *k*-mers and builds no index. With constant-time rank and select, finding the defects, including the walks and the sweep over the dead nodes, and Part 2 of Algorithm 1 take *O*(*n*) time. Each source is first spelled out with *k* select queries. Part 1 then places the nodes of its dummy chain with *O*(*k*) rank queries, and the at most |*Q* | (*k−* 1) new nodes can be sorted by position with a radix sort in *O*(*n* + |*Q*| *k*) time. In total, the dummy nodes are repaired in *O*(*n* + |*Q*| *k*) time, against the *O*(*nk*) algorithm of our earlier work [9]. The group starts and the merge plan of two inputs still take *O*(*nk*) time (Section 2), so a whole operation does too.

### 3.2 Colors for every operation

Section 3.1 in combination with the merge plan gives the SBWT of the result of every operation: the extraction by colors, the intersection, the difference and the union. A colored index also stores the distinct color sets and the color set identifier of every key *k*-mer (Section 2), so every operation must decide which color set each *k*-mer of the result has, and which *k*-mers are key *k*-mers.

The color sets follow from the operation. The union gives every *k*-mer *x* of the result the color set col_*A*_(*x*) *∪* col_*B*_(*x*), where *x* may be in one input only, and the difference gives it col_*A*_(*x*). The extraction by colors and the intersection have two modes. By default, they keep only the colors that the operation selects: the extraction of a set *C* of colors gives *x* the color set col(*x*) *∩ C*, and the intersection gives it col_*A*_(*x*) *∩* col_*B*_(*x*), which may be empty. When they *keep all colors*, they give *x* its full color set, col(*x*) or col_*A*_(*x*) *∪* col_*B*_(*x*), which tells, for every kept *k*-mer, which other colors share it.

#### 3.2.1 Extracting by colors

Given a set *C* of colors, the extraction by colors keeps the *k*-mers whose color set contains a color of *C*, in either mode. In the example, removing c, that is, keeping *C* = *{a, b, z}* , keeps every *k*-mer except `TCCT`, `CTGT` and `TGCT`, which occur in c alone. To achieve this, the extraction by colors finds the kept *k*-mers in one walk over the unitigs of the input, which tests every colored subunitig once, since all its *k*-mers have the same color set. The extraction of Section 3.1 then computes the SBWT of the result (Figure 1b).

Subsequently, the key *k*-mers of the result have to be chosen, that is, the last *k*-mer of every colored subunitig of the result (Section 2). Some of them end a unitig of the result, and are found by walking its unitigs: in the example, these are `GCTG, CTTG, GTTG and TGCA`. The others are the *k*-mers after which the color set changes inside a unitig of the result: in the example, `TTGC`, with *{b, z}* , before `TGCA`, with *{b}*. Since the color set of a *k*-mer in the result only depends on its color set in the input, the color set also changes there in the input, so such a *k*-mer ends a colored subunitig of the input, as `TTGC` does. To find them, the walk over the input subunitigs records the last *k*-mer of every kept colored subunitig, and Part 2 of Algorithm 1 returns its node in the result: in Table 2, `TTGC` is node 4 of the input and node 5 of the result.

Not every recorded *k*-mer is needed, however. `CGCT` ends a unitig of the input, because `GCTG` also has the in-neighbor `TGCT`, but in the result `TGCT` is gone, and `CGCT` runs into `GCTG`, both with *{a}*. To avoid unnecessary key *k*-mers, the walk gives every distinct color set a fingerprint of the colors it *keeps*, the XOR of a random 128-bit value per color [38], and records the last *k*-mer of a colored subunitig only where this fingerprint changes. Two sets of kept colors have the same fingerprint with probability 2^*−*128^, since the XOR of the random values of the colors in which they differ is uniform. Only then is a needed key *k*-mer missed.

In summary, after loading the input, the extraction by colors runs in five steps. The *walk* computes the group starts of the input, walks its unitigs, marks the kept *k*-mers, and records the last *k*-mer of every kept colored subunitig. The *SBWT* step extracts the SBWT of the result (Section 3.1). The *key* step marks the recorded *k*-mers, the last *k*-mer of every unitig of the result, and new checkpoints every *d k*-mers. The *look-up* step finds the color set of every marked *k*-mer in the input, and chooses one to represent every distinct color set. The *store* step looks up these color sets again, and stores each of them once [38].

#### 3.2.2 Intersection, difference and union

The intersection, the difference and the union work in a similar way to the extraction by colors, but with the merge plan in place of the input. The intersection and the difference read *K* off the merge plan and apply the extraction to it (Section 3.1), and the union keeps the whole merge plan. Besides the unitig ends of the result, their key *k*-mers come from walking the colored subunitigs of the inputs whose colors the result uses: both inputs for the union and the intersection, and *A* for the difference. This finds every color change of the intersection and the difference for the same reason as for the extraction by colors: their *k*-mers are in every input whose colors they use, so where the color set changes in the result, it also changes between the same two *k*-mers in one of these inputs. Since the color set of a *k*-mer in the result can depend on both inputs, a fingerprint of the colors of one input cannot tell whether it changes, so, unlike the extraction by colors, these operations keep every *k*-mer that they record, including a few that change no color set. The color set of a key *k*-mer combines its color sets in the inputs that have it, as the operation prescribes.

The union needs one more rule, because where the color set changes, an input may have only one of the two *k*-mers. If it has only the *k*-mer before the change, this *k*-mer ends a unitig of the input, since its only out-neighbor in the union is missing there, so it is marked as before. If it has only the *k*-mer after the change, the change may go unnoticed. For example, let *A* have `ACGT` and `CGTA`, both with *{a}* , and *B* have `CGTA` and `GTAC`, both with *{b}*. In the union, the color set changes after ,`ACGT`, from *{a}* to *{a, b}* , so `ACGT` must be a key *k*-mer. Neither input marks it, however: in *A*, `ACGT` and `CGTA` have the same color set, and *B* does not have `ACGT`. The change is visible in *B* from the other side, however: `CGTA`, the *k*-mer after the change, has no in-neighbor in *B*, so a colored subunitig of *B* starts there. The union therefore also marks the in-neighbors, in the union, of the first *k*-mer of every colored subunitig of its inputs, here `ACGT`.

##### Observation 3

*The union marks every k-mer after which the color set changes inside a unitig of the union*.

Let *y* follow *x* inside a unitig of the union, with a different color set. Then col_*I*_ (*x*) ≠col_*I*_ (*y*) in one input *I*, where a *k*-mer that is not in *I* has the empty set. Inside a unitig, *y* is the only out-neighbor of *x*, and *x* the only in-neighbor of *y*. If *I* has both, they are thus consecutive in a unitig of *I*, and *x* ends a colored subunitig of *I*. If *I* has only *x*, then *x* ends a unitig of *I*. If *I* has only *y*, then *y* starts a colored subunitig of *I*, and the union marks its in-neighbor *x*. In all three cases, *x* is marked.

## 4 Results

Section 4.1 compares editing with rebuilding the index from the genomes, Section 4.2 compares insertion with the three-way merge, and Section 4.3 applies the operations to AllTheBacteria.

### Setup

All experiments ran on a shared server with two Xeon Gold 6430 processors (64 cores, without hyper-threading), 503 GiB of RAM and a local NVMe SSD. We used 32 threads unless stated otherwise (thread scaling: Figure A1). We ran our implementation^3^ and GGCAT 2.2.0 [39]. All indexes have *k* = 31. The datasets are 3 682 *E. coli* assemblies [40] (341 M *k*-mers, 108 colors per *k*-mer on average) and 121 034 *S. pneumoniae* assemblies from AllTheBacteria (414 M *k*-mers, 1 199 colors per *k*-mer, a 43 GB index). We report medians of 5 runs on *E. coli* and 3 on *S. pneumoniae* (1 for the two largest rebuilds).

### Baseline

We compare extraction and merge, a union that adds genomes to an index, with a *rebuild* : building their result from scratch, from the genomes that it contains. For this we use the fastest pipeline that we found, in three steps: GGCAT computes the unitigs of the genomes without colors, `sbwt build` builds the SBWT of these unitigs, and Themisto [2, 38] adds the colors to this SBWT, reading the genomes again. For results of at most 100 genomes, we also ran Themisto alone, which builds the SBWT itself, and report it when it is faster. The time of a rebuild is the sum of its steps, and its memory the largest of their peaks. A rebuild needs the genomes, while editing does not. Intersections and differences have no such baseline, as their results index no set of genomes.

### 4.1 Editing vs. rebuilding

The time of an extraction depended mostly on the input index, while that of a rebuild depended on the result (Figure 2). On *E. coli*, an extraction took 18 to 54 s whatever was kept, while a rebuild grew from 4 s for one genome to 13 min for all but one. Editing was faster as soon as the result had more than about 100 genomes, or about 3% of the collection, and 14 times faster for removing one genome, or 100 (Tables 3 and A1). On *S. pneumoniae*, removing 1, 10, 100 or 1 000 genomes took 17 min in every case, and removing one genome was 17 times faster than a rebuild, which read the other 121 033 genomes (80 GB compressed) and took 4.7 h. There, rebuilding was faster for up to at least 1 000 genomes, if they are at hand.

**Table 3.** Editing an index vs. rebuilding the result from the genomes. Time in m:ss or h:mm:ss, peak memory in GiB. The speedup is the time of the rebuild divided by that of the edit, so editing is faster above 1*×*. Figure 2 shows all extractions (details in Table A1 in the supplement). A merge adds genomes to the index of all other genomes, and its time is that of building the index of the added genomes plus that of the merge.

| dataset | edit | edit |  | rebuild |  | speedup |
|---|---|---|---|---|---|---|
|  |  | time | peak | time | peak |  |
| *E. coli* | extract: remove 1 genome | 0:54 | 4.1 | 13:01 | 5.3 | 14× |
|  | merge: add 1 genome | 0:04 + 0:58 | 4.1 | 13:24 | 5.2 | 13× |
|  | merge: add 10% (368) | 1:11 + 1:03 | 4.1 | 13:24 | 5.2 | 6× |
| *S. pneumoniae* | extract: remove 1 genome | 16:57 | 80.0 | 4:44:27 | 43.3 | 17× |
|  | merge: add 1 genome | 0:02 + 15:46 | 80.0 | 4:37:29 | 43.6 | 18× |

**Figure 2.**
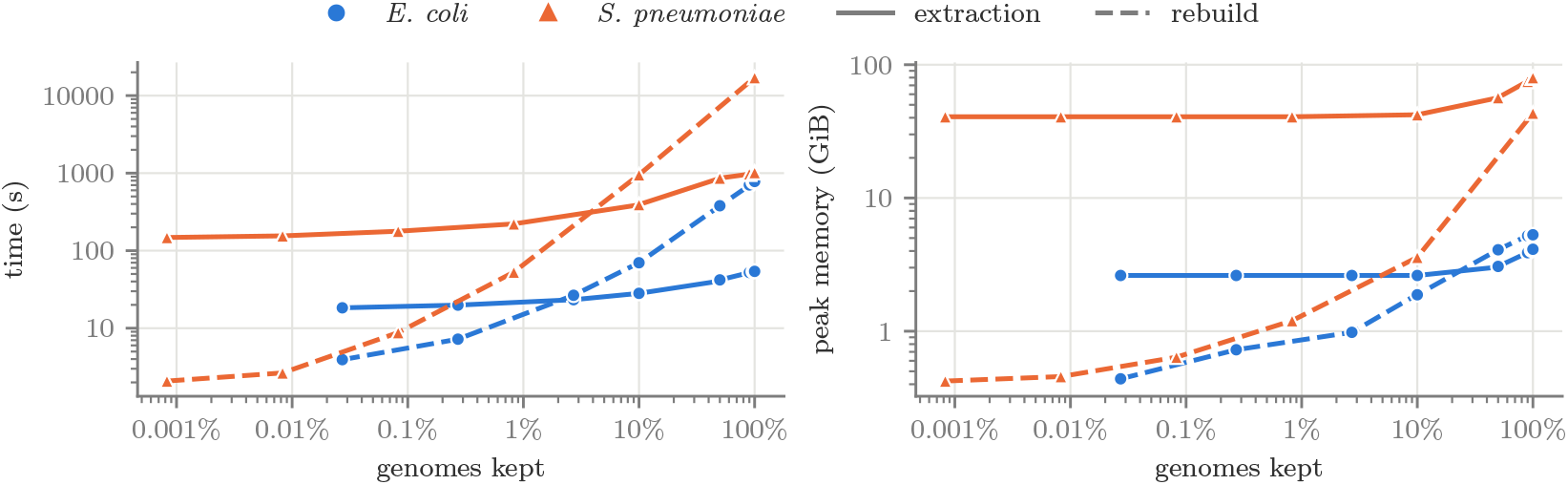
Extraction vs. rebuild from the genomes, against the fraction of the genomes kept, on *E. coli* (3 682 genomes) and *S. pneumoniae* (121 034 genomes), on log scales.

On *S. pneumoniae*, 80% of the time of removing one genome went into the look-up and store steps of Section 3.2.1, whose cost grows with the number of colors that they read: the distinct color sets of *S. pneumoniae* take 42 GB against 1.7 GB for *E. coli*, so removing one genome took 19 times as long, with only 1.2 times as many *k*-mers (Table A2 in the supplement). The SBWT step, which contains the extraction of Section 3.1, took only 1.5 to 4.1 s: under 10% of the time on *E. coli* and 2% on *S. pneumoniae*.

An edit loads its input index, so even keeping one genome of *S. pneumoniae* took 41 GiB, the size of the index (rebuild: 0.4 GiB); with the result on top, removing one took 80 GiB (rebuild: 43 GiB). *E. coli* took 2.6 to 4.1 GiB, less than a large rebuild. Editing thus saves time and the genomes, at the cost of memory.

The typical use of a merge is to add new genomes to an existing index. Its cost was set by the size of its result: merging two *E. coli* indexes into the full index took 58 to 66 s, whether the second index had 1, 36, 368 or 1 841 genomes. Adding genomes therefore cost building their index plus about one minute, 13 times faster than a rebuild for one genome, and 6 times for 368 genomes. On *S. pneumoniae*, adding one genome took 946 s, 18 times faster than rebuilding the full index, and, as for extraction, most of this time went into the color sets. The supplement covers merging many indexes, and indexes that share genomes (Section A.4).

### 4.2 Repairing the SBWT

The intersection and the difference of two indexes are extractions (Section 3.1), and we used them to compare insertion (Section 3.1.2) with the three-way merge of our earlier work [9]^4^. Section 3.1 predicts that insertion takes *O* (*n* + |*Q*| *k*) time instead of *O* (*nk*). We ran both repairs on 31 operations: differences and intersections of *E. coli* indexes that are nested, overlap or are disjoint, extractions by colors, and three on *S. pneumoniae*. Table 4 shows five that span the range of repair times, and Table A3 in the supplement lists them all.

**Table 4.**
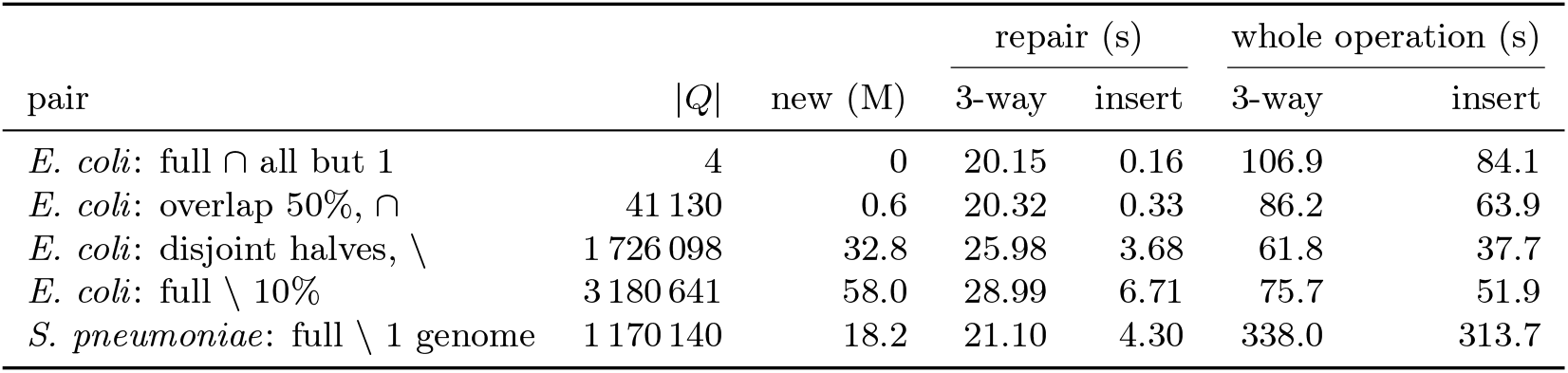
Repairing the SBWT in a difference (\) or an intersection (*∩* ): the three-way merge of our earlier work [9] (3-way) against insertion (insert, Section 3.1.2). In every pair, the two inputs together hold all *k*-mers of the full index, so their merge plan has about as many positions as the full index has nodes: *n* = 343 million on *E. coli* and 418 million on *S. pneumoniae*. |*Q*| : sources; new: dummy nodes inserted (0 when the dummy chains of all sources already exist in an input). Repair: from finding the sources until the merge plan of the result is complete; whole operation: wall time of the command. Intersections keep the intersection of the color sets.

|  |  |  | repair (s) |  | whole operation (s) |  |
|---|---|---|---|---|---|---|
| pair | \|*Q*\| | new (M) | 3-way | insert | 3-way | insert |
| *E. coli*: full ∩ all but 1 | 4 | 0 | 20.15 | 0.16 | 106.9 | 84.1 |
| *E. coli*: overlap 50%, ∩ | 41 130 | 0.6 | 20.32 | 0.33 | 86.2 | 63.9 |
| *E. coli*: disjoint halves, \ | 1 726 098 | 32.8 | 25.98 | 3.68 | 61.8 | 37.7 |
| *E. coli*: full \ 10% | 3 180 641 | 58.0 | 28.99 | 6.71 | 75.7 | 51.9 |
| *S. pneumoniae*: full \ 1 genome | 1 170 140 | 18.2 | 21.10 | 4.30 | 338.0 | 313.7 |

On the pairs of Table 4, insertion repaired the SBWT 4 to 126 times faster than the three-way merge. As predicted, the three-way merge took 20 s even for 4 sources, since its merge plan makes *k* passes over all positions, while the time of insertion followed |*Q*| : with over a million sources, most of it went into spelling out the sources and placing their dummy chains, the |*Q*| *k* term of its bound. Both used similar peak memory. The rest of the time, mostly the merge plan of the two inputs and the color sets, does not depend on the repair, and was about the same for both. The whole operation thus became 1.3 to 1.6 times faster on *E. coli*, and 1.1 times on *S. pneumoniae*, where the color sets dominate.

#### Correctness

Every result that we could compare with construction was equivalent to it, with the same *k*-mers, the same color set for every *k*-mer, and the same color names. This covers the extractions and merges of Sections 4.1 and A.4, and the intersections of the full *E. coli* index with a part of it, which must give that part. For every pair *A, B* of an intersection and a difference, we also checked that |*A \ B*| + |*A ∩ B* |= *A* , and that both color modes keep the same *k*-mers. Insertion and the three-way merge also gave identical indexes in all 31 operations run with both.

### 4.3 Case study: AllTheBacteria

The species index of AllTheBacteria [6] (release 2024-08) has one color per species call, including 80 mixed calls of two species and the color ‘unknown’ for the 73 159 samples without a call. It contains 192 billion *k*-mers of 12 733 species, and takes 194 GB on disk. Building it from the unitigs of AllTheBacteria took 15 h (11 h for the SBWT and 4 h for the colors), using up to 455 GiB of memory. Its color of *M. tuberculosis* is suspiciously large. *M. tuberculosis* is a clonal species with little horizontal gene transfer, whose genomes differ by at most a few thousand single-nucleotide variants [41], and a single-nucleotide variant adds at most 2*k* new *k*-mers, in both orientations. Even the 180 094 genomes of its color should therefore add little to the 8.8 million *k*-mers of one genome, yet the color has 4.35 billion *k*-mers, as many as about 495 genomes.

Extracting the *M. tuberculosis* color from the species index, with all colors kept, gave an index *X* of every *k*-mer of the color, together with every other color that has it. It took 3 h 34 min, mostly to compute the group starts of the index and walk its unitigs, at a peak of 386 GiB, about twice its size on disk, as every bit vector over its *n* nodes takes 22 GiB. Every further question was then answered on *X* and on smaller indexes, in 4 to 17 min and at most 36 GiB per step (Table A4 in the supplement). An index of the same genomes, built from their assemblies, was cheaper, as Section 4.1 predicts for a small result: 1 h 59 min at 17.3 GiB (Section A.7). It records only the *M. tuberculosis* genomes, however, while the extraction also tells, for every *k*-mer of the color, which of the other 12 732 species share it, and this is what the rest of this section uses.

From *X* we extracted the color itself, *M* , and the *k*-mers of *M* that also occur in another color, *O*. The color ‘unknown’ and the mixed calls contain samples of *M. tuberculosis* themselves, so we also extracted the *k*-mers shared with the colors of single species, *O*_sp_. Two small indexes serve as references: *R*, of the reference genomes *M. tuberculosis* H37Rv and *M. bovis* AF2122/97, and *H*, of the human genome CHM13 [42].

Only 12.8% of *M* is shared with another species, mainly common contaminants and species isolated together with *M. tuberculosis*, led by *A. baumannii, E. coli* and *E. faecalis*, and non-tuberculous mycobacteria. In the metadata of AllTheBacteria, sylph [43] also detects them in many *M. tuberculosis* samples, for example *E. coli* in 1 387, *M. abscessus* in 1 087 and *P. aeruginosa* in 903. Another 8.6% is shared only with ‘unknown’ and the mixed calls, and contains about 70% of *R*. The remaining 78.6% occurs in no other color. Its long unitigs are mostly low in GC, with 69% of their *k*-mers below 60% GC, against 65.6% for *M. tuberculosis*, which fits organisms that have no color of their own. Its short unitigs resemble *M. tuberculosis* and are likely variants and assembly errors. Human sequence is 0.3% of *M* (Table A5 in the supplement).

Set operations and one heuristic filter then removed most of the candidate foreign sequence. First, *M* ^*′*^ = (*M \ O*_sp_) \ *H* drops the *k*-mers shared with other species and the human ones. Simulated reads of 8 foreign species had no hits in *M* ^*′*^, unlike H37Rv reads. Second, we select the unitigs of ((*M \ O*) \ *H*) \ *R* with at least 301 *k*-mers and less than 60% GC, two heuristic thresholds, and build an index *F* from them, the only step that is not a set operation. Then *M* ^*′′*^ = (M′*\ F* ) *∪* (*M ∩ R*) removes them, and puts back any part of the reference genomes that a difference removed. *M* ^*′′*^ kept 66.2% of *M* and all of *R*.

The index that we built from the assemblies covers the *M. tuberculosis* genomes of release 2024-08 and of the next release, 2025-05, with one color per group of genomes. Extracting its genomes of release 2024-08 gave exactly the *k*-mers of *M* , which confirms the extraction at the scale of AllTheBacteria. Release 2025-05 revised the species calls, and drops 31.6% of the *k*-mers of *M* , about as many as our cleaning (33.8%), but only 36% of these are also cleaned by us. Instead, the new release moves whole genomes out of the color, mostly noisy assemblies of *M. tuberculosis*, which a color of the species index cannot single out. It keeps most of what our species index flags as possibly foreign, 613 million *k*-mers of *F* and 362 million of *O*_sp_. Set operations thus found, without the assemblies, contamination that the next release keeps (Section A.7).

## 5 Conclusion

Colored *k*-mer indexes need not be rebuilt every time the collection they represent changes. We have shown that genomes can be removed from or extracted from a colored SBWT index, and two such indexes combined by union, intersection or difference, without the genomes, with a result equivalent to construction.

Editing complements construction rather than replacing it: the cost of an edit is set by its inputs and that of a rebuild by its result, so editing is the method of choice when the result is large, as when a few genomes are removed, and rebuilding when the result is small and the genomes are at hand. Editing also keeps what a rebuild discards: an extracted species can retain, for every *k*-mer, all other species that share it, which let us trace contamination in AllTheBacteria with set operations and one heuristic filter.

In indexes with many colors, color sets take most of the time of an edit, so they are the main target for speedups. To make editing easy to adopt, a natural next step is to host a server that keeps the index of AllTheBacteria and its group starts in memory, so that users can obtain an index of any species or clade without downloading either the genomes or the full index.

## A Supplementary material

### A.1 Extractions and merges

**Table A1:** Extractions and merges of Section 4.1: editing an index vs. rebuilding the result from the genomes. Time in m:ss or h:mm:ss, peak memory in GiB. The speedup is the time of the rebuild divided by that of the edit, so editing is faster above 1*×*. The extractions are ordered by the fraction of the genomes kept, as in Figure 2. A merge adds genomes to the index of all other genomes, and its time is that of building the index of the added genomes plus that of the merge (i.e., we assume one index is already built and must be updated). The rebuilds of all or all but one genome of *S. pneumoniae* are single runs.

| dataset | edit | edit |  | rebuild |  | speedup |
|---|---|---|---|---|---|---|
|  |  | time | peak | time | peak |  |
| *E. coli* | extract: keep 1 genome | 0:18 | 2.6 | 0:04 | 0.4 | 0.2× |
|  | extract: keep 10% (368) | 0:28 | 2.6 | 1:10 | 1.9 | 2.5× |
|  | extract: keep 50% (1 841) | 0:42 | 3.1 | 6:20 | 4.1 | 9× |
|  | extract: remove 100 genomes | 0:54 | 4.1 | 12:49 | 5.2 | 14× |
|  | extract: remove 1 genome | 0:54 | 4.1 | 13:01 | 5.3 | 14× |
|  | merge: add 1 genome | 0:04 + 0:58 | 4.1 | 13:24 | 5.2 | 13× |
|  | merge: add 10% (368) | 1:11 + 1:03 | 4.1 | 13:24 | 5.2 | 6× |
| *S. pneumoniae* | extract: keep 1 genome | 2:28 | 40.7 | 0:02 | 0.4 | 0.01× |
|  | extract: keep 10% (12 103) | 6:33 | 42.1 | 15:53 | 3.6 | 2.4× |
|  | extract: remove 1 genome | 16:57 | 80.0 | 4:44:27 | 43.3 | 17× |
|  | merge: add 1 genome | 0:02 + 15:46 | 80.0 | 4:37:29 | 43.6 | 18× |

### A.2 Time per step of the extraction by colors

**Table A2:** Time per step of the extraction by colors, in m:ss, for the extractions of Table A1 (medians of 5 runs on *E. coli* and 3 on *S. pneumoniae*, 32 threads). *Load* : reading the input index from disk. *Walk, SBWT, key, look-up* and *store*: the steps of Section 3.2.1. *Write*: writing the result to disk.

| dataset | extraction | load | walk | SBWT | key | look-up | store | write | total |
|---|---|---|---|---|---|---|---|---|---|
| *E. coli* | keep 1 genome | 0:02.6 | 0:12.9 | 0:01.5 | 0:00.4 | 0:00.8 | 0:00.3 | 0:00.0 | 0:18.3 |
|  | keep 50% (1 841) | 0:02.6 | 0:13.9 | 0:03.0 | 0:05.4 | 0:10.8 | 0:05.6 | 0:00.8 | 0:42.0 |
|  | remove 1 genome | 0:02.6 | 0:13.7 | 0:02.9 | 0:06.7 | 0:17.9 | 0:09.0 | 0:01.8 | 0:54.1 |
| *S. pneumoniae* | keep 1 genome | 0:55.6 | 1:17.7 | 0:02.2 | 0:00.4 | 0:09.0 | 0:02.4 | 0:00.0 | 2:27.9 |
|  | keep 10% (12 103) | 0:54.8 | 1:28.0 | 0:03.1 | 0:02.2 | 2:05.7 | 1:55.5 | 0:02.0 | 6:33.1 |
|  | remove 1 genome | 0:55.9 | 1:20.8 | 0:04.1 | 0:08.6 | 8:22.2 | 5:13.5 | 0:52.9 | 16:57.1 |

*S. pneumoniae* has 1.2 times as many *k*-mers and key *k*-mers as *E. coli*, but its distinct color sets have 26 435 colors on average against 1 027, and take 42 GB against 1.7 GB. The look-up and store steps, whose cost grows with the number of colors that they read, took 30 times as long.

### A.3 Thread scaling

Keeping half of *E. coli* took 675 s on one thread and 42 s on 32, 16 times faster. Every step but loading and writing the index, which take about 3 s, sped up with the number of threads (Figure A1).

**Figure A1:**
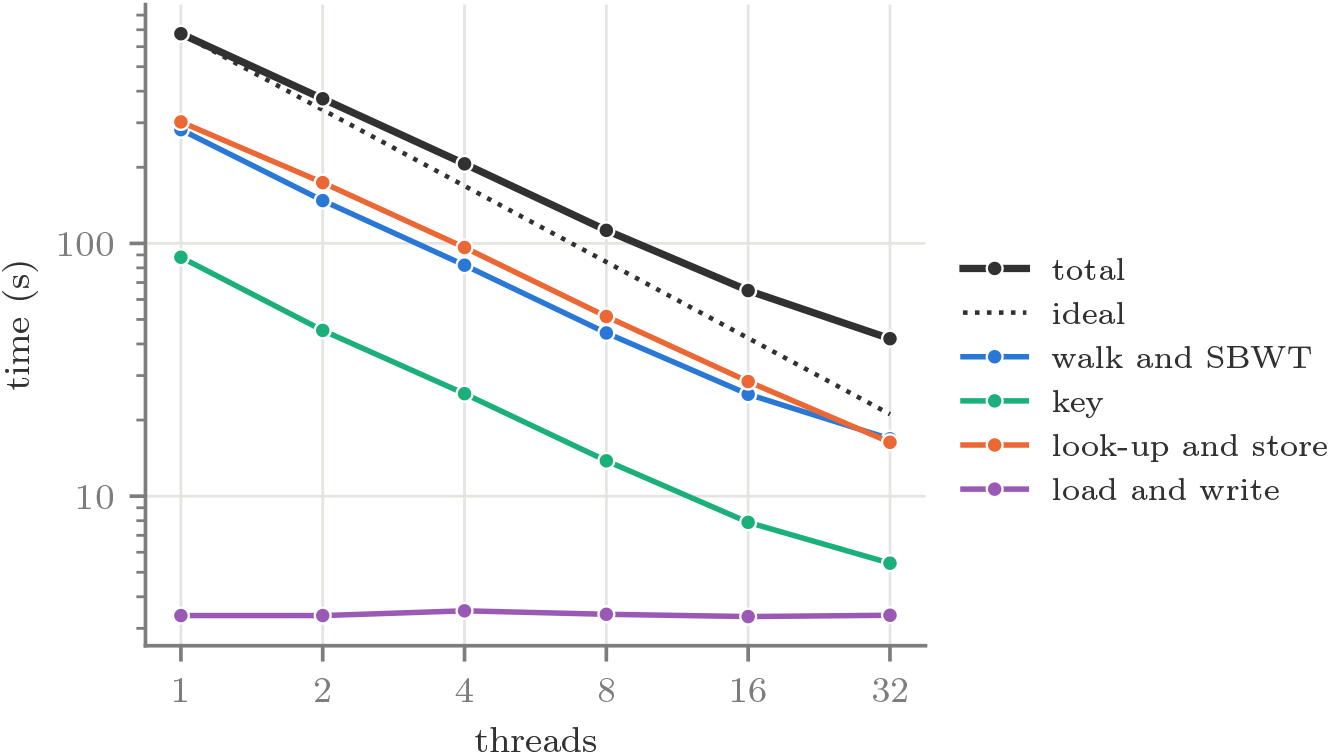
Time per step of keeping half of *E. coli* (1 841 genomes) against the number of threads, log scales, medians of 5 runs. The steps are grouped as in Table A2. The dotted line is the time on one thread divided by the number of threads.

### A.4 Merging many indexes, and indexes that share genomes

To combine many indexes, merging them in pairs, in a balanced tree, is faster than adding them one at a time to a growing index, since each genome then goes through log_2_ of the number of indexes merges instead of up to that number: for *E. coli* split into 4, 8 or 16 equal parts, the tree took 142, 246 and 379 s, against 158, 318 and 617 s one at a time in a linear fashion. Inputs may also share genomes, for example two releases of a collection. Their colors are then matched by name, and each shared genome appears once in the result. When the two *E. coli* indexes share 10, 50 or 100% of the genomes, so that their merge is again the full index, the merge took 68, 75 and 84 s, against 66 s for two disjoint halves, as the inputs grow with the shared genomes. On *S. pneumoniae*, merging two halves took 940 s, close to the 946 s of adding one genome to the other 121 033.

### A.5 Intersections and differences

Table A3 lists all 31 operations on which we compared the two repairs of Section 4.2. On *E. coli*, the pairs answer different questions about the genomes. The intersection of the full index with one genome, keeping all colors, gave the 10.3 million *k*-mers of the genome together with every genome that shares each of them, in 1.55 million distinct color sets, while the intersection of the color sets left only the genome itself as color. The difference of the full index and all genomes but one gave the 5 226 *k*-mers that occur in the remaining genome alone, and the difference of the full index and one genome gave the 331.0 million *k*-mers that this genome lacks. The intersection of two disjoint halves, keeping all colors, gave the 183.0 million *k*-mers that occur in both halves, and their difference the 93.8 million that occur in the first half only. The difference of half of the genomes and the full index was empty. Insertion was faster in every operation that repairs anything, and both repairs gave identical indexes in all 31. When no source misses its dummy chain, as when removing one genome by colors, neither repairs anything, and the two take about the same time.

**Table A3:** All operations on which we compared the three-way merge of our earlier work [9] (3-way) with insertion (insert), as in Table 4. Medians of 5 runs on *E. coli* and 3 on *S. pneumoniae*. Pair: the inputs *A* and *B*, where full is the full index, a percentage the index of that fraction of the genomes, and overlap the two indexes that share that fraction of the genomes. Colors: the intersection of the color sets (*∩* ), or all colors kept (all). Result: *k*-mers of the result, in millions. *n*: the positions of the merge plan of the two inputs for a difference or an intersection, as in Table 4, and the nodes of the single input, the full index, for an extraction by colors. It is 343 million on *E. coli* and 418 million on *S. pneumoniae* in every operation. |*Q*|, new, repair and whole operation as in Table 4.

|  |  |  |  |  | repair (s) |  | whole operation (s) |  |
|---|---|---|---|---|---|---|---|---|
| pair | colors | result (M) | \|*Q*\| | new (M) | 3-way | insert | 3-way | insert |
| *E. coli*, difference *A* \ *B* |  |  |  |  |  |  |  |  |
| full, 1 genome |  | 331.0 | 1 479 495 | 27.1 | 19.80 | 3.87 | 68.8 | 52.5 |
| full, 10% |  | 210.9 | 3 180 641 | 58.0 | 28.99 | 6.71 | 75.7 | 51.9 |
| full, 50% |  | 64.6 | 1 584 686 | 30.2 | 26.04 | 3.38 | 63.4 | 38.5 |
| full, all but 1 |  | <0.1 | 139 | <0.1 | 20.19 | 0.17 | 51.2 | 28.9 |
| overlap 10% |  | 88.1 | 1 593 122 | 30.4 | 25.30 | 3.42 | 60.9 | 37.2 |
| overlap 50% |  | 45.6 | 847 323 | 16.6 | 23.20 | 2.05 | 56.1 | 32.8 |
| disjoint halves |  | 93.8 | 1 726 098 | 32.8 | 25.98 | 3.68 | 61.8 | 37.7 |
| 50%, full |  | 0 | 5 719 | 0 | 19.99 | 0.22 | 47.7 | 26.3 |
| *E. coli*, intersection *A* ∩ *B* |  |  |  |  |  |  |  |  |
| full, 1 genome | ∩ | 10.3 | 140 | 0 | 14.90 | 0.23 | 42.2 | 25.9 |
| full, 1 genome | all | 10.3 | 140 | 0 | 14.96 | 0.26 | 54.5 | 38.7 |
| full, 10% | ∩ | 130.3 | 6 214 | 0 | 18.32 | 0.29 | 64.2 | 44.1 |
| full, 10% | all | 130.3 | 6 214 | 0 | 18.29 | 0.29 | 79.7 | 60.6 |
| full, 50% | ∩ | 276.7 | 13 499 | 0 | 20.00 | 0.27 | 87.3 | 64.5 |
| full, 50% | all | 276.7 | 13 499 | 0 | 20.12 | 0.26 | 95.6 | 73.6 |
| full, all but 1 | ∩ | 341.3 | 4 | 0 | 20.15 | 0.16 | 106.9 | 84.1 |
| full, all but 1 | all | 341.3 | 4 | 0 | 20.12 | 0.16 | 106.8 | 83.8 |
| overlap 10% | ∩ | 197.3 | 148 769 | 2.9 | 20.44 | 0.63 | 70.7 | 48.1 |
| overlap 10% | all | 197.3 | 148 769 | 2.9 | 20.20 | 0.62 | 87.2 | 64.7 |
| overlap 50% | ∩ | 266.0 | 41 130 | 0.6 | 20.32 | 0.33 | 86.2 | 63.9 |
| overlap 50% | all | 266.0 | 41 130 | 0.6 | 20.25 | 0.34 | 96.8 | 73.4 |
| disjoint halves | all | 183.0 | 204 818 | 4.1 | 20.34 | 0.77 | 84.6 | 62.0 |
| 50%, full | ∩ | 276.7 | 13 499 | 0 | 20.09 | 0.26 | 88.2 | 64.7 |
| 50%, full | all | 276.7 | 13 499 | 0 | 19.99 | 0.26 | 95.4 | 74.1 |
| *E. coli*, extraction by colors |  |  |  |  |  |  |  |  |
| keep 1 genome |  | 10.6 | 206 | <0.1 | 12.87 | 0.19 | 30.8 | 18.3 |
| keep 10% |  | 129.0 | 20 964 | 0.3 | 13.04 | 0.28 | 41.2 | 28.2 |
| keep 50% |  | 276.7 | 30 573 | 0.1 | 13.00 | 0.24 | 55.0 | 42.0 |
| remove 100 genomes |  | 338.9 | 8 149 | <0.1 | 11.94 | 0.18 | 66.0 | 53.8 |
| remove 1 genome |  | 341.3 | 0 | 0 | 0.00 | 0.00 | 54.9 | 54.1 |
| *S. pneumoniae* |  |  |  |  |  |  |  |  |
| full \ 1 genome |  | 410.0 | 1 170 140 | 18.2 | 21.10 | 4.30 | 338.0 | 313.7 |
| keep 10% |  | 127.0 | 22 612 | <0.1 | 16.37 | 0.30 | 434.4 | 393.1 |
| remove 1 genome |  | 414.1 | 0 | 0 | 0.00 | 0.00 | 1025.8 | 1017.0 |

### A.6 The steps of the AllTheBacteria case study

Only the two extractions from the species index need hours and hundreds of GiB. Every other step of Section 4.3 takes at most 17 min and 36 GiB (Table A4).

**Table A4:** Time (m:ss or h:mm:ss) and peak memory (GiB) of the steps of the case study (Section 4.3), one run each, 32 threads. Only the first two read the species index of AllTheBacteria.

| result | operation | time | peak |
|---|---|---|---|
| *X* | extraction of *M. tuberculosis* from the species index, keeping all colors | 3:33:41 | 386.3 |
| *M* | extraction of *M. tuberculosis* from the species index | 3:33:59 | 386.0 |
| *M* | extraction of *M. tuberculosis* from *X* | 5:53 | 16.9 |
| *O* | extraction of all other colors from *X* | 7:17 | 25.7 |
| *M* \ *O* | difference | 10:39 | 29.4 |
| *O*<sub>sp</sub> | extraction of the colors of single species from *X* | 6:09 | 23.2 |
| *M* \ *O*<sub>sp</sub> | difference | 10:55 | 27.1 |
| (*M* \ *O*) ∩ *H* | intersection | 11:14 | 29.8 |
| *M* ∩ *R* | intersection | 4:31 | 14.8 |
| *M*′ | difference | 16:35 | 36.2 |
| *F* | build from its unitigs | 8:20 | 6.5 |
| *M*″ | merge | 8:17 | 11.0 |

### A.7 The *M. tuberculosis* color of AllTheBacteria in detail

We count *k*-mers as the index stores them, in both orientations, unless we say canonical *k*-mers.

**Table A5:** The *M. tuberculosis* color *M* of the species index of AllTheBacteria: where its *k*-mers come from (top, as disjoint parts that add up to |*M*|), and what remains after cleaning (bottom).

|  | *k*-mers | share of *M* |
|---|---|---|
| shared with a single species (*O*<sub>sp</sub>) | 555 170 492 | 12.8% |
| shared only with ‘unknown’ or a mixed call (*O* \ *O*<sub>sp</sub>) | 376 083 438 | 8.6% |
| only in *M*, human or reference ((*M* \ *O*) ∩ (*H* ∪ *R*)) | 12 573 202 | 0.3% |
| only in *M*, the rest, in unitigs of > 300 *k*-mers | 1 307 765 636 | 30.0% |
| in unitigs of 32 to 300 *k*-mers | 1 377 037 322 | 31.6% |
| in unitigs of ≤ 31 *k*-mers | 724 015 264 | 16.6% |
| *M* | 4 352 645 354 | 100% |
| *M*′ = (*M* \ *O*<sub>sp</sub>) \ *H* | 3 781 891 694 | 86.9% |
| *F*, long low-GC unitigs removed from *M*′ | 904 165 174 | 20.8% |
| *M*″ = (*M*′ \ *F*) ∪ (*M* ∩ *R*) | 2 880 434 636 | 66.2% |

#### What the color contains

The species that share the most with *M* are led by *A. baumannii* (35.7 M canonical *k*-mers), *E. coli, E. faecalis, P. aeruginosa* and *S. epidermidis*, followed by non-tuberculous my-cobacteria such as *M. abscessus* and *M. avium. O* contains almost all of the reference genomes, with only 82 of their *k*-mers in *M O*. Among the *k*-mers that occur in no other color, those of short unitigs have only 27% of their *k*-mers below 60% GC. These unitigs form 460 115 bubbles, 58% of them a single substitution, and many tips and isolated *k*-mers. What *M* ^*′′*^ holds beyond one genome is mostly short and medium sequence with characteristics like *M. tuberculosis*, the variation and errors of 180 094 genomes.

#### The index built from the assemblies

Of the genomes that are *M. tuberculosis* in release 2024-08 or 2025-05, 172 536 are so in both (*K*), 7 558 only in 2024-08 (*D*), and 15 822 only in 2025-05 (*N* ). We built an index of these 195 916 genomes with one color per group. For each of the 51 archives of AllTheBacteriathat contain them (3.5 GB to download in total, the largest unpacking to 17.7 GB of assemblies), GGCAT computed the unitigs of each group and Themisto built an index from them, and the 51 indexes were then merged in a balanced tree. This took 1 h 59 min at a peak of 17.3 GiB (62 min for the 51 indexes, 56 min for the merges), and gave an index *KDN* of 4.62 billion *k*-mers in 3.96 GB. Extracting the colors *K* and *D* from *KDN* gave exactly the *k*-mers of *M*.

#### Comparison with the next releas

Of the *k*-mers of *M* , the new release drops 31.6% and our cleaning removes 33.8%, and only 36% of the *k*-mers that the new release drops are also removed by the cleaning. The new release moves whole genomes out of the color: *D* is 4.2% of the genomes of the color but holds 31.6% of its *k*-mers, about 182 000 of its own per genome, and 64% of these are in *M* ^*′′*^, so these genomes were mostly noisy assemblies of *M. tuberculosis*, which a color of the species index cannot single out, since it does not record which genome a *k*-mer comes from. The cleaning instead removes the sequence that the index identifies as foreign, which the new release largely keeps: it drops about a third of *F* (32.2%) and of *O*_sp_ (34.8%), barely more than of *M* ^*′′*^ (30.4%), so that 613 million *k*-mers of *F* and 362 million of *O*_sp_ remain in its color. On human sequence the two agree: the new release drops 72.6% of the human *k*-mers of *M* , and the cleaning all but the 12 that are also in the reference genomes. Together, the cleaning and the curation of the new release keep 2.0 billion of the 4.35 billion *k*-mers of *M*.

## Acknowledgements

This work has benefited from funding from the French State under the France 2030 program, reference ANR-21-IDES-0006. The European Metropolis of Lille and the University of Lille are also acknowledged for their funding and support of the project WILL-CHAIRES-25-001-BOSSA.

## Footnotes

1 https://github.com/LoreDepuydt/colored-sbwt-setops

2 https://crates.io/crates/sbwt

3 https://github.com/LoreDepuydt/colored-sbwt-setops/tree/recomb27; the scripts of all experiments, with how to run them, are in its folder benchmarks.

4 Ported into our implementation: https://github.com/LoreDepuydt/colored-sbwt-setops/tree/recomb27-threeway

